# Noncoding RNAs endogenously rule the cancerous regulatory realm while proteins govern the normal

**DOI:** 10.1101/791970

**Authors:** Anyou Wang, Rong Hai

## Abstract

All cancers share a commonality in genome activation regulated by a systems endogenous network distinct from normal tissue, but such a network remains elusive. Here, we unearth a systems regulatory network endogenous for all types of cancers and normal human respectively from massive data, including all RNAseq data available from SRA and TCGA, and reveal distinctive systems realm for cancer and normal. In the cancerous realm, noncoding RNAs, especially pseudogenes, dominate endogenous network modules and centrality, and they work as the strongest systems inducers that cis-regulate their targets. However in the normal realm proteins dominate the entire endogenous network centrally controlled by ribosomal proteins and they trans-regulate their targets. Our finding establishes a systems picture of an endogenous mechanism overlooking the cancerous and normal realm, in which noncoding RNAs rule the overall cancer realm while proteins govern the normal one. This fundamentally refreshes the conventional concept of cancerous mechanism.

## INTRODUCTION

All cancers generally result from endogenous genomic abnormality distinct from normal human tissue[1–4]. Understanding the universal regulatory mechanisms endogenous in all cancers helps to develop the general therapeutic strategy to combat cancers.

Numerous gene regulations and networks have been identified for tumorigenesis, but most of these gene interactions are specific for a given cancer type, which refers to a specific organ and tissue[5–9]. Thus tumorigenesis mechanisms have been mostly marked as cancer type specific. However, certain regulations have been found endogenously in all cancer types. For example, TP53 has been typically characterized as an universal suppressor endogenous for all cancers[5]. More recently, a pseudogene PTENP1 has also been identified to be an endogenous regulator that regulates PTEN in examined cancer types[8,10]. Given million gene regulations in human genome, the magnitude of endogenous cancerous regulations for all cancer types should be very large. These gene regulations usually assemble a systems cancerous regulatory network distinct from that of normal human. Revealing such systems networks helps to advance our deep insights toward fundamental mechanisms of cancer and normal human physiology.

Uncovering such endogenous regulatory networks faces challenges. First of all, the human regulatory network is complex, and emerging noncoding RNAs complicate this network[8,9,11]. In addition, numerous conditions and factors contribute to a regulatory network. Secondly, we have not developed any appropriate biological approaches to reveal a real natural gene regulation. Current biological approaches like knockout suffer several limitations such as transcript compensation and genome alteration[12]. Knocking out a single gene normally results in alterations of thousand gene activation, leading to biased picture of gene regulations. Thirdly, current computational inferences suffer high noise, with low accuracy <50%[13,14]. Due to data limitation, computational inferences are also heavily biased to the investigated data set. Therefore, the current knowledge of genome-wide gene regulations still looks like the picture derived from blind men and an elephant. A complete picture of any regulatory realm still remains elusive.

In this present study, we developed a software, FINET[14], to infer the precise endogenous gene regulations. Technically, FINET optimizes elastic-net and stability-selection and infers any gene regulation without any presumption, such as a noncoding RNA to a protein_coding gene or vice versa. With optimizing parameters and filtering out the condition-dependent interactions, FINET actually selects endogenous gene regulations that were consistently true in all conditions. These gene regulations reasonably assemble an universal network for a given dataset. To maximize the universality and to make our inferences endogenous for all cancers and normal human tissue, we processed massive data, including all human RNAseq data available from Sequence Read Archive (SRA 274469 samples) and The Cancer Genome Atlas (TCGA 11574 samples). These data contained all conditions and cancer types, which promised us to infer endogenous regulations. After inferring endogenous networks from these SRA and TCGA data respectively for normal human and cancers, we generated quantitative patterns from these networks to reveal endogenous rulers for the cancerous and normal tissue.

## RESULTS

### Systems regulatory networks endogenous in cancers and normal human

To assemble a systems regulatory network endogenous for all conditions in all cancers and normal human respectively, we first needed a complete set of data representing endogenous genomic activation universal for all conditions. SRA and TCGA provided such data. SRA RNAseq contained various data sets of all conditions. A gene regulation that is extracted from SRA 274469 samples and is consistently true in all SRA conditions represents an endogenous regulation universal for normal human(**Figure S1**, materials and methods). These endogenous regulations in systems level assemble a systems network endogenous for normal human. Similarly, TGCA provided RNAseq data for 11574 samples containing 32 cancer types[16] and systems gene regulations extracted from these cancerous data constructed a systems endogenous network for cancers.

Secondly, we need a software that can infer an accurate unbiased gene regulation. We developed algorithms and a software, FINET[14]. Compared to the current software with high noise during inference, FINET significantly improves the accuracy, and it can efficiently and accurately infer unbiased gene interactions with >94% precision, true positives/(true positives+false positives) (materials and methods). FINET filters out all condition-dependent interactions and only keeps the true endogenous ones independent from any conditions such as biological sample heterogeneity and sequencing technique variations.

We employed FINET to search all possible gene regulations in human genome by systematically treating each gene as a target and selecting its regulators from the rest of all annotated genes(**Figure 1A**, materials and methods). In this way, each gene has an equal chance to be a target or a regulator without any presumption, regardless of its gene category in protein_coding or noncoding RNA. This search was separately performed for SRA and TCGA data, and the result eventually was assembled into a network for normal human and cancer respectively. The normal network contained 19,721 nodes (genes) and 63,878 edges (interactions), and the cancerous one included 25,402 nodes and 61,772 edges (**Fig. 1B-1C**).

**Figure 1.**
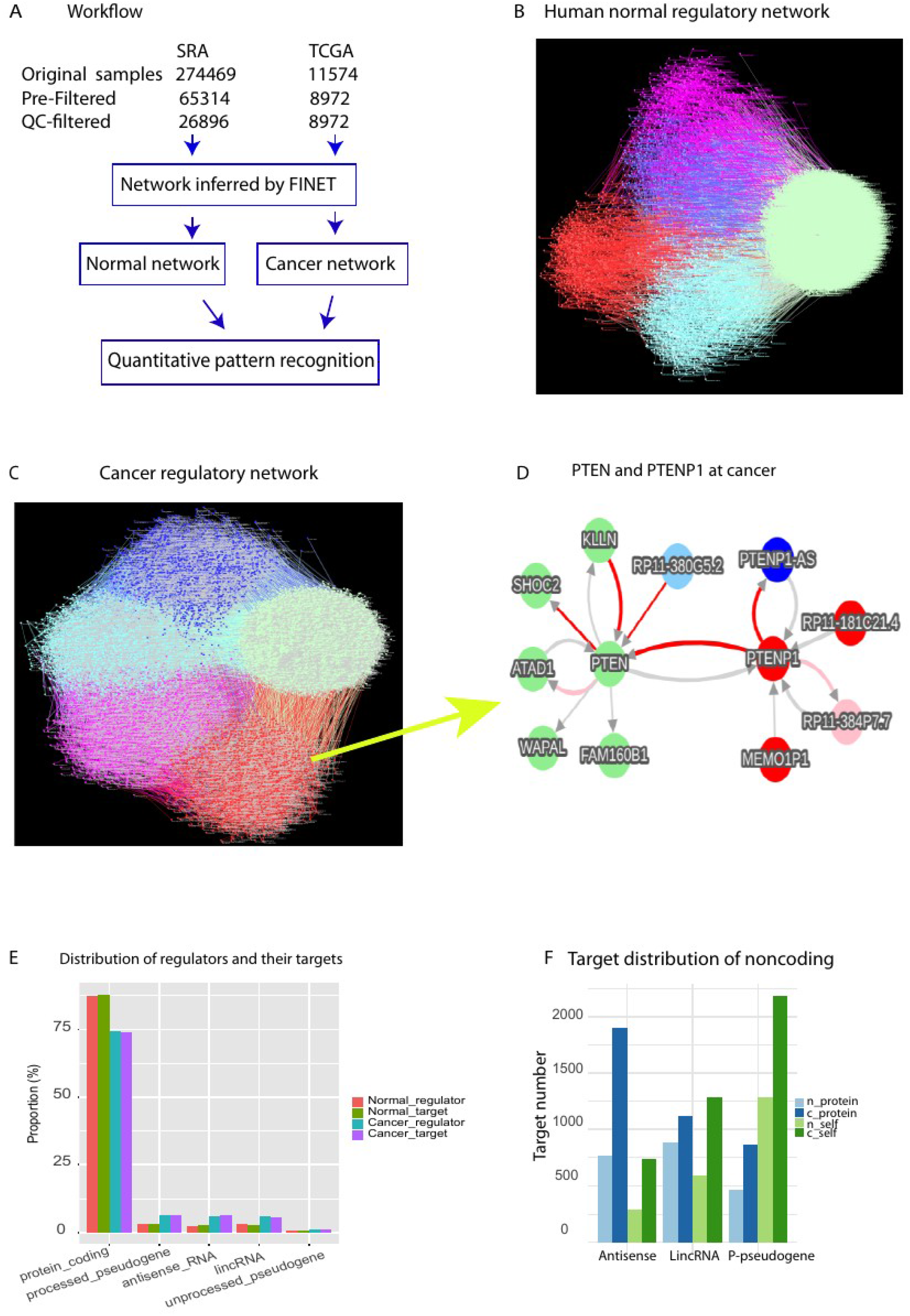
Gene regulatory networks endogenous in cancers and normal human. A, the workflow of this study. B-C, Completed gene regulatory network endogenous in normal (B) and cancer(C). The nodes (genes) and edges (interactions) were grouped into 5 gene category sets, including protein (light green), antisense(blue), lincRNA (pink), p_pseudogene (red), and the rest (other, lightblue). D, an example of sub_network, PTEN interacting with PTENP1 in the cancer network directly extracted from our network database. E, overall distribution of regulators and their targets at normal and cancer. The top 5 most abundant categories were shown. F, The target distribution of three categorized noncoding RNAs, antisense, lincRNA, and p-pseudogene, at normal (n_) and cancer (c_). Targets were counted separately when the three individual noncoding RNAs as regulators. Self denotes the targets as self-categorized genes. For example, c_self antisense represents cancerous antisense RNAs that were targeted by cancerous antisense RNAs.

As expected, our networks were much less complex than those of current reports because we only collected the reliable interactions endogenous in all conditions, yet these two networks actually represented two distinct regulatory realms endogenous for normal human and cancer at systems level, in which all endogenous layered crosstalk of any pair of genes were included. As a validation, our network contained an interaction between PTEN (protein_coding) and PTENP1 (a pseudgene of PTEN) (**Figure 1D**), and this interaction only existed in the cancer network but did not exist in the normal one[17], consistent to experimental reports showing it only in cancers[8,10]. This indicated our network with high reliability and specificity. Furthermore, in contrast to conventional approaches showing PTENP1 as a regulator for PTEN only, our systems network expands this PTENP1-PTEN interaction to a cancerous PTENP1 regulatory network including several novel PTENP1 interactions (**Figure 1D**). PTENP1 also interacted with its-own antisense_RNA (PTENP1_AS), two pseudogenes (RP11-181C21.4, MEMO1P1), and a lincRNA(RP11-384P7.7). These natural endogenous interactions provide a complete systems regulatory picture for PTENP1. Moreover, because our network is universally true for all types of cancers, our result also suggested PTENP1 regulating PTEN as a universal endogenous regulation in all cancers although it was uncovered by conventional approaches with limited cancer types and conditions.

Similarly, a complete systems regulatory picture of any universal endogenous regulation can be easily extracted from our network online[17]. Strikingly, an antisense RNA RP11-335k5.2, which was recently uncovered by our clinical data analysis as the most strongest inducer for all cancers[16], was consistently found here in our cancer network[17], but not in the normal network. This indicated RP11-335k5.2 indeed as an endogenous cancer driver for all types of cancers.

Overall, our networks provide a reliable and comprehensive resource for understanding the complete systems pictures of endogenous regulations in the cancer and normal genome.

### Overall noncoding RNA crosstalk are unexpectedly activated at cancers

Multilayered crosstalk among proteins and various types of noncoding RNAs play key roles in physiologic states but the complete picture of crosstalk endogenous in cancers and normal human tissues remains elusive[8,9]. Here we first examined the picture by grouping activated genes into gene sets via set algorithms[18], which clusters a network into sub-network sets on the basis of node and edge properties. By using gene annotated categories as node attributes, we separated the entire network into 5 gene category sets, including protein_coding (referred as protein hereafter), lincRNA, processed-pseudogene (p-pseudogene), antisense RNA (antisense), and others that pooled the rest of gene categories**(Figure 1B-1C)**.

In normal tissue, the majority of proteins and p-pseudogenes were mostly either separated or self-targeted, in which targets and their regulators at the same gene category(referred as self-regulation thereafter), but most of antisense RNAs and lincRNAs were highly crosstalked to proteins(**Figure 1B**). However, in cancer these 5 sets were overall separated, and the density of the protein set became less than normal(**Figure 1B_1C**), indicating that cancerous protein-protein crosstalk declined but crosstalk within noncoding RNAs increased. Statistically, we counted the regulators and their targets in each gene category in both normal and cancer (**Table S1, Table S2**). Overall, at normal the total crosstalk around proteins occupied 87.7% (56039/63878), and the rest 12.3% was around noncoding RNAs (**Table S1**). However, for cancer the overall crosstalk around proteins significantly declined to 73.9% (45660/61774), and crosstalk around noncoding RNAs increased to 26.1% (**Table S2**) (p-value = 0.02157, Pearson’s Chi-squared test with Yates’ continuity correction, referred as chisq-test thereafter). We next counted the specific gene category interactions. The interactions from proteins to proteins at normal counted for 82.5% (52692/63878, **Table S1**), but declined to 64.8% at cancer (40053/61774, **Table S2**). The protein regulators and protein targets also decreased from normal to cancer, but noncoding RNA regulators and targets dramatically increased in cancer (**Figure 1E**, p-value < 2.2e-16, Chisq-test). These indicated the primary regulatory crosstalk shifted from normal protein domination to cancerous noncoding RNAs.

To further explore the detailed targets of noncoding RNAs, we plotted the primary targets of three abundantly categorized noncoding RNAs, including antisense, lincRNA, and p-pseudogene. Targets of noncoding RNAs primarily contained not only proteins but also self-regulated genes such as p-pseudogenes primarily regulate p-pseudogenes (**Figure 1F, Table S1, Table S2**). These self-targets of all three categorized noncoding RNAs were significantly induced by cancer (p-value = 1.46e-08, **Figure 1F**). For example, p-pseudogenes targeted self-targets, p-pseudogenes, with significantly increasing from 1288 at normal to 2185 at cancer**(Figure 1F**). This indicated that noncoding RNAs, especially p-pseudogenes, increase self-regulation in cancer.

Together, protein crosstalk dominate the normal network, but noncoding RNA crosstalk become unexpectedly activated in cancers. Noncoding RNAs significantly turn to self-regulation in cancers.

### Network module composition shifts from normal proteins to cancerous noncoding RNAs

To understand the module differences between normal and cancer network, we examined module member compositions. We identified modules by network topology[19] and then clustered modules into either protein module (proteins occupied > 50% of members in a module) or noncoding module (noncoding RNAs > 50% of members in a module, materials and methods). Modules with 50% of proteins or noncoding RNAs were ignored. At normal protein modules occupied 60.52% out of total 38 modules and noncoding modules only took 28.94% (**Figure 2A, table S3**)), while cancerous modules significantly changed their compositions, in which protein modules reduced to 47.29% and noncoding modules increased to 45.94% of total 74 cancer modules (p-value = 0.02963, chisq-test)( **Figure 2A**, **table S4**). Theoretically the network modules execute the primary functions for a network. This module pattern shifting from proteins to noncoding RNAs suggested noncoding RNAs as the key rulers in cancer regulatory realm.

**Figure 2.**
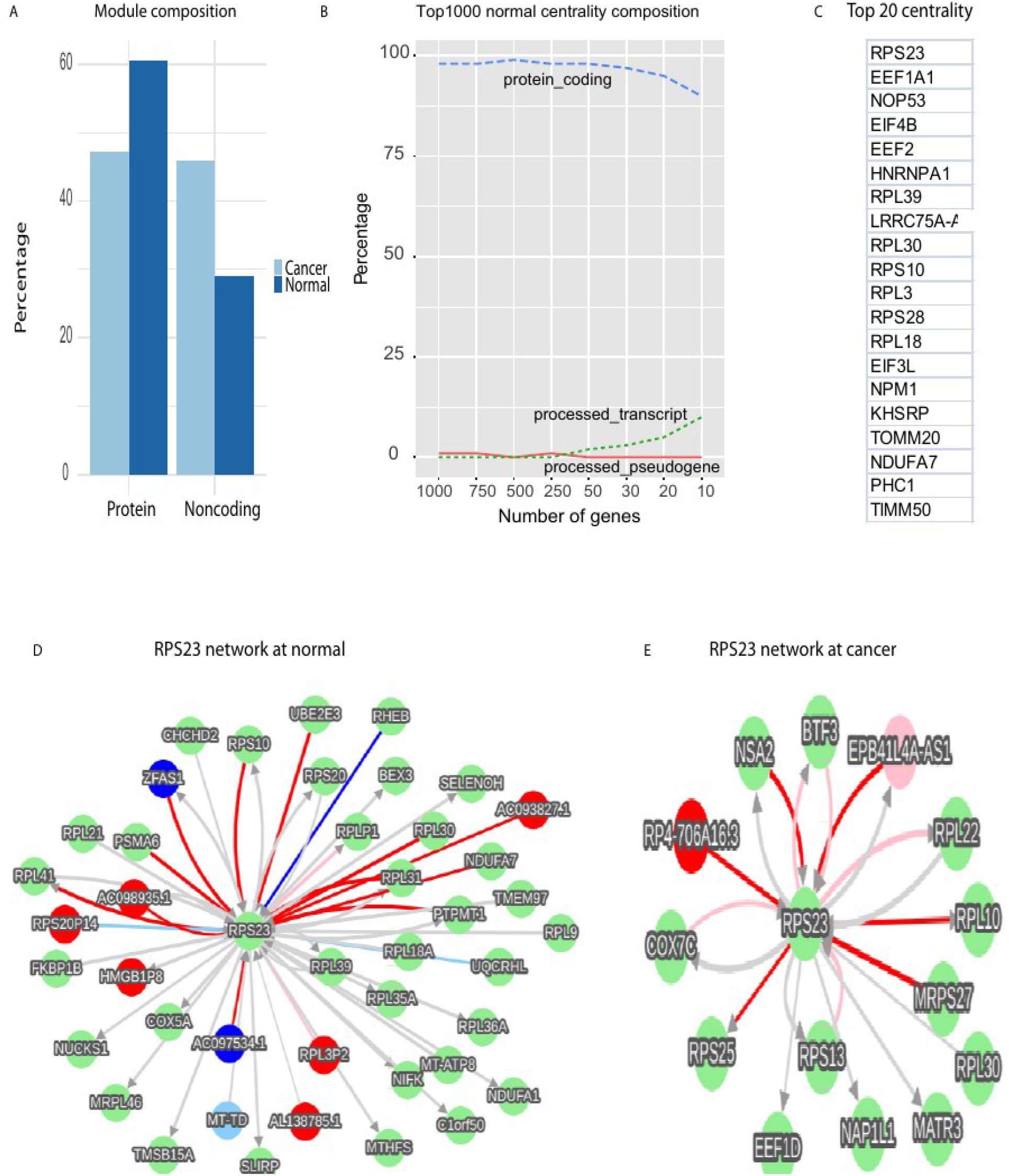
Network module compositions and normal network centrality. A, module composition differences between normal and cancer network. B, Compositions of top 1000 normal network centrality. C, Top 20 normal centrality. D-E, RPS23 first neighbors in normal (D) and cancer(E).

### Noncoding RNAs serve as the centrality in cancerous network but ribosomal proteins dominate in the normal

To understand the core controllers of the normal and cancerous networks, we investigated the centrality of normal and cancer networks (materials and methods). At normal proteins worked as the primary centrality (top 1000, **Figure 2B**) and ribosomal proteins dominated the top 20 centrality in normal (**Fig. 2C**). The top 1 centrality, RPS23, abundantly interacted with proteins and noncoding RNAs **(Figure 2D),** but at cancer the interactions of RPS23 declined dramatically (**Figure 2E**). Based on gene ontology (GO)[20], these top 20 centrality in normal networks performed crucial functions in translation (RPL18, RPL3, RPL30, RPL39, RPS10, RPS23, RPS28). Consistently, the functions for the whole normal network and modules (**table S3**) were also relevant to translation and negative regulations, suggesting ribosomal proteins as the delicately regulatory core of the normal human genome.

In contrast, p-pseudogenes dominated the cancerous centrality (**Figure 3A**, materials and methods) and most of these centrality worked as cancer inducers (regulators with coefficient > 0 and < 0 were respectively referred as inducers and repressors during FINET inferences, materials and methods, **Figure 3B**). Most of these inducers were p-pseudogenes(**Figure 3C**), and all top 20 centrality were p-pseudogenes (**Figure 3D**). This indicated p-pseudogenes as the primary rulers for cancers.

**Figure 3.**
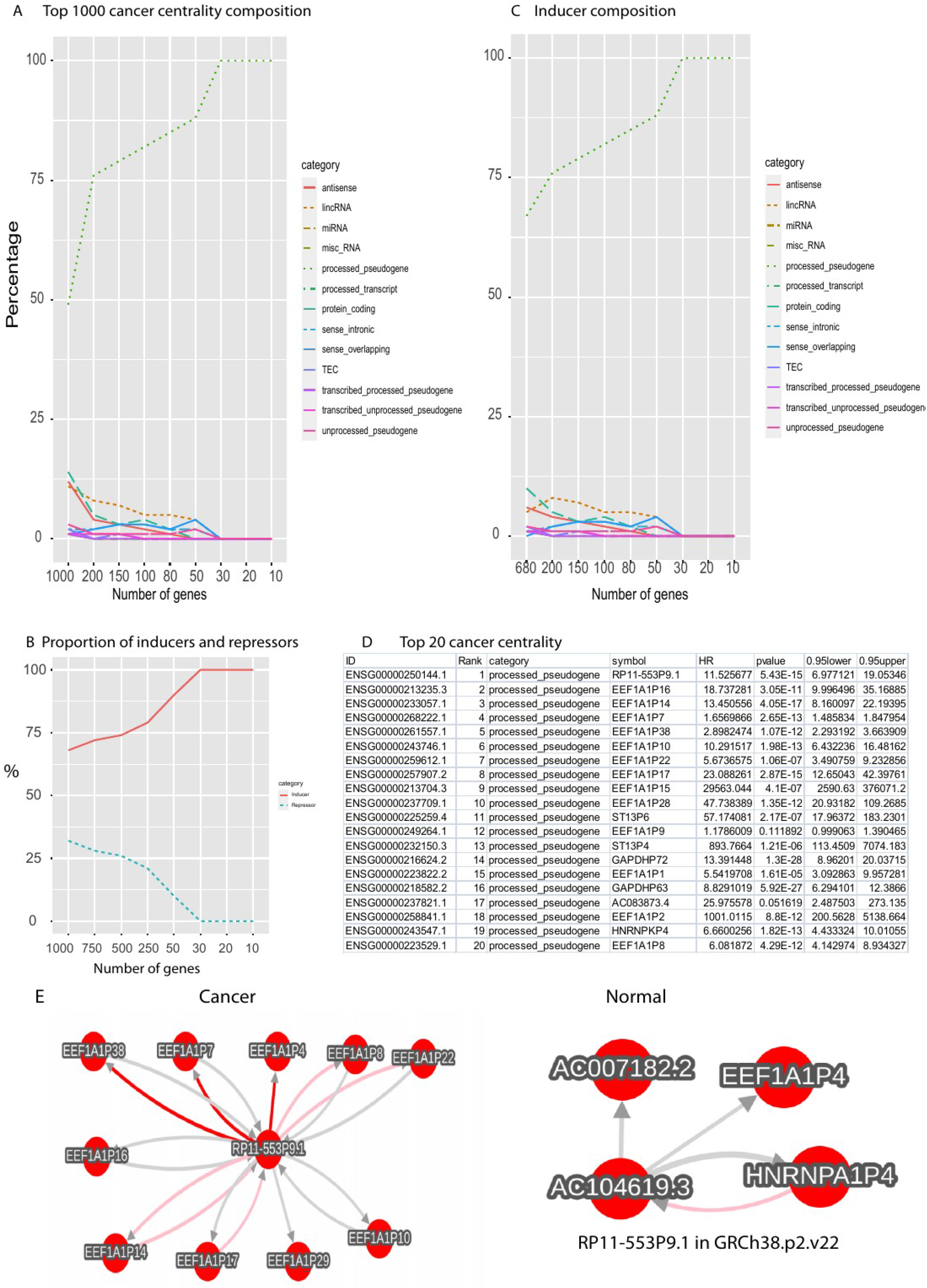
Cancer network centrality. A, Compositions of top 1000 cancer network centrality. B, proportion of inducers and repressors in top 1000 cancer network centrality. C, compositions of inducers in top cancer centrality(B). D, top 20 cancer centrality. E, ENSG00000250144.1 has more interactions in cancers than normal. Please note that the gene symbol was different in the two annotation versions as labeled in the figure and our database.

These pseudo-gene interactions escalated in cancers have been shown in our database. For example, cancers activated much more interactions in the top listed pseudo-gene(**Figure 3D**), ENSG00000250144.1, than normal (**Figure 3E**).

These data suggest that ribosomal proteins serve as the most important regulatory core for protein-dominated normal network, but noncoding RNAs, especially p-pseudogenes, primarily control the center of cancerous realm.

### Noncoding RNAs and proteins respectively serve as the strongest regulators in the cancerous and normal network

To understand the strongest regulators governing normal and cancer genome and to make our pattern robust, we examined the composition of the top 300 regulators and their corresponding targets based on their absolute coefficient rankings. For pattern recognition and clear illustration, we only presented any gene category with abundance > 10%. At normal, proteins worked as the strongest inducers. From the top 300 to top 10 inducers, proteins occupied 60% to 50% respectively (**Figure 4A left**). LincRNAs came next and occupied ~20%. These inducers mostly targeted proteins and p-pseudogenes (**Figure 4A right**). Yet in cancer, proteins even did not show up (<10%), instead, noncoding RNAs dominated the top inducers, including p-pseudogene, antisense RNA and lincRNA (**Figure 4B left**). For example, p-pseudogenes counted 70% out of top 10, suggesting p-pseudogenes as the primary strongest drivers in cancer genome, instead of proteins as conventionally thought. Interestingly, these cancerous inducers almost purely targeted proteins(**Figure 4B right**). This suggested that proteins work as targets at cancer instead of as cancerous drivers. The conventional practice treating protein-coding genes as cancerous drivers is very misleading. Consistently, our result from big clinical data also found p-pseudogenes as the primary drivers universal for all types of cancers[16].

**Figure 4.**
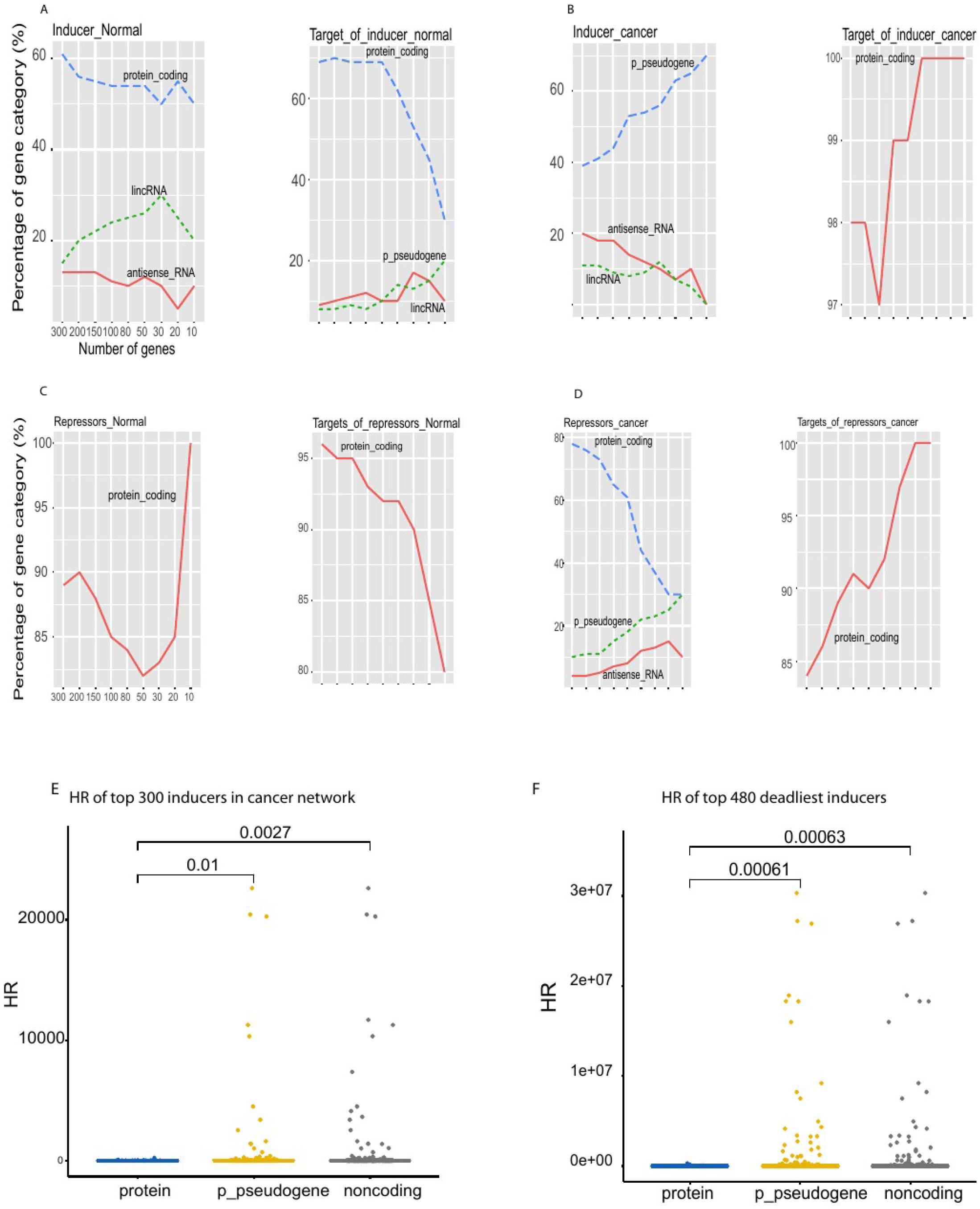
Top 300 strongest inducers and repressors at the normal and cancer network. The composition of strongest inducers and repressors at normal and cancer network (A-D). A, The top 300 strongest inducers (left) and their targets (right) of normal network. B, the top 300 strongest cancerous inducers and their targets. C, the top 300 strongest repressors and their targets at normal. D, The top 300 strongest cancerous repressors and their targets. Clinic data of top cancerous inducers(E-F). E, comparison of hazard ratio (HR) between protein, p-pseudogenes and noncoding RNAs in top 300 strongest inducers in the cancer network built by this present study. F, HR profiling of top 480 deadliest inducers directly extracted from Cox proportional-hazards model analysis of all TCGA RNAseq data**[16]**. P-values (above line) were calculated by t-test.

As for the strongest repressors, almost all repressors and their targets were proteins at normal (**Figure 4C**). However, cancerous repressors contained proteins, p-pseudogenes, and antisense RNAs, with at least 10% at each (**Figure 4D**). Surprisingly, regardless of normal and cancerous repressors, almost all their targets were proteins >85%, and noncoding RNA targets in any categories were too low to show (<10%). This pattern revealing proteins as targets for both inducers and repressors in cancer interprets why the current observations have focused on proteins, yet treating protein-targets as cancerous drivers is fundamentally misleading. Nonconding RNAs, especially p-pseudogenes, serve as the primary universal drivers for all types of cancers.

### Noncoding RNAs serve as the deadliest inducers in cancers

To understand the cancerous association of noncoding RNAs, we calculated the HR (hazard ratios) of top 300 inducers in the cancer network as described above (Materials and methods). Among these inducers, p-pseudogenes and all noncoding RNAs (including p-pseudogenes) had significant higher HR than proteins, with pvalue=0.01 between p-pseudogenes and proteins and pvalue=0.0027 between noncoding RNAs and proteins (**Figure 4E**). To further confirm this result, we examined these HR differences between proteins, p-pseudogenes and noncoding RNAs in top deadliest inducers derived from unbiased survival analysis of all cancer type data from TCGA[16](Materials and methods). Similarly, both p-pseudogenes and noncoding RNAs had significant higher HR than proteins, with pvalue=0.00061 and 0.00063 respectively(**Figure 4F**). These results consistently indicated that noncoding RNAs, especially p-pseudogenes, play more important roles in causing cancer death than proteins. This and our previous results[16] provide strong systems evidences to validate our systems network results showing noncoding RNAs as the most important drivers for tumorigenesis, instead of proteins.

### Noncoding RNAs primarily turn to regulate their local targets at cancer

Understanding the systems distribution of distances between regulators and their targets help to understand the functional framework of genome regulations but it remains debated[8,9,21–23]. To capture the systems profiling of target distances altered by cancer, we compared it to that of normal in top four gene categories, including protein, p-pseudogene, lincRNA, and antisense. To overlook the profiling, we clustered the targets by chromosomes via using set-algorithm as done in figure 1 above. At normal, all chromosome sets were mixed up but these sets were clearly separated at cancer (**Figure 5A-5B**), indicating that noncoding RNAs increasingly targeted their self-chromosome targets at cancer compared to normal. Statistically, most of normal genes worked as trans-regulators regulating their targets outside their chromosomes(**Figure 5C**). Especially, more than 80% p-pseudogene targets located outside chromosome, and 70% protein and 55% lincRNA targets were also located outside chromosome. Furthermore, p-pseudogenes and proteins rarely regulated their targets with overlapped sequences (inside genes, **Figure 5C**). However, in cancer most regulators of all categories turn to regulate their local targets (<1M bp **Figure 5D**). Specifically, more than 80% of lincRNAs and antisense RNAs worked locally.

**Figure 5.**
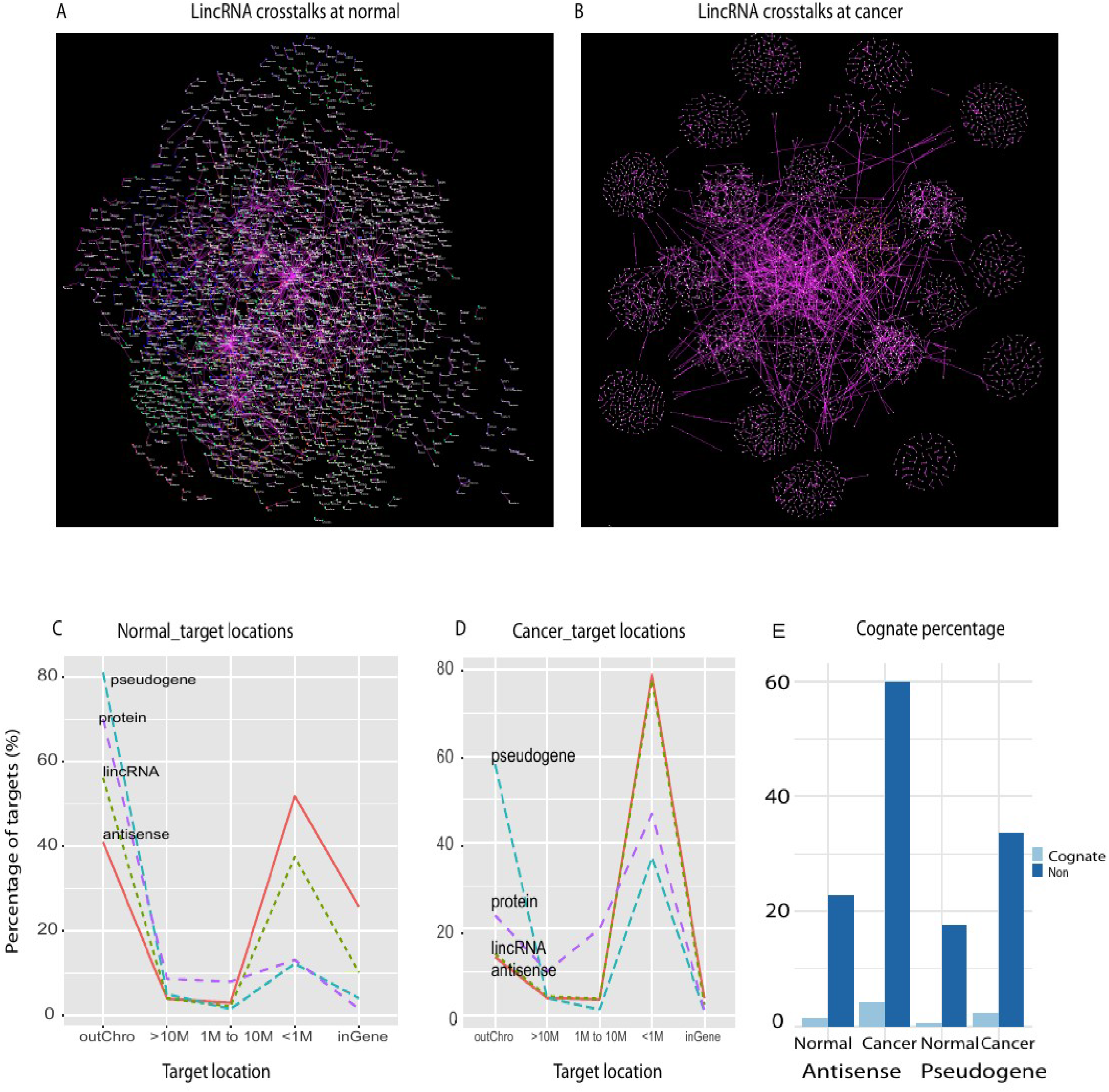
Target distance distribution. A-B, lincRNA regulatory network in normal (A) and cancer(B). These two networks were grouped by chromosome to show crosstalk between chromosomes. C-D, target location distribution of top abundant gene categories (>10%) in normal (C) and cancer (D). OutChro represents targets locating outside chromosome, and M denotes million bp inside the chromosome. E, the percentage of cognates and non-cognates targeted by antisense RNAs and pseudogenes at normal and cancer. Non denotes non-cognate

However, these antisense RNAs and pseudogenes rarely regulated their cognates at both normal and cancer(**Figure 5E**). Furthermore, cancer stimulated the non-cognate proportion. The non-cognate rate for antisense increased from normal 22.8% to cancerous 60%, and this for p-pseudogene also increased from 17.6% (normal) to 33.5% (cancer). In contrast, the cognate proportion shifted slightly from 85 (1.4% normal) to 241 (4.2% cancer) for antisense, and from 73 (0.6% normal) to 254 (2.3% cancer) for pseudo-gene (**Figure 5E**). This suggested primary noncoding RNAs as cis-regulators in cancers, but not as cognate regulators as recently proposed[24].

This together suggests that regulations switch from normal trans-regulations to cancerous cis-regulations.

## DISCUSSION

This study revealed a complete systems picture of endogenous regulatory mechanisms regulating the cancer and normal realm, in which noncoding RNAs endogenously rule the cancerous regulatory realm while proteins govern the normal. Numerous regulatory mechanisms have been uncovered for regulating cancers and normal physiology, but they are biased to a given biological experiment and are condition-dependent and thus are not universally endogenous for all conditions. The systems mechanism endogenous across all conditions remain unknown. Here, we revealed that proteins control normal human regulatory realm at systems level and ribosomal proteins endogenously govern the core of the normal realm. Ribosomal proteins have been known as important factors in controlling cell type specific physiology and pathology[25], but we found more important role for them in which they actually work as an universal endogenous center to regulate whole human normal realm via interacting with other proteins and noncoding RNAs. This realm is dominated by proteins working as trans-regulators to regulate proteins as their primary targets, consistent with current practices in biology in which proteins are treated as both key regulators and targets. However, this normal protein-dominant realm cannot be applied to cancers. Cancers are endogenously regulated by noncoding RNAs. Noncoding RNAs, especially p-pseudogenes, serve as the primary centrality and the strongest inducers, and they also control the cancerous modules functioning for the entire systems realm. This parallels with our recent observation from clinical data showing noncoding RNAs as the universal deadliest drivers for all types of cancers[16]. Our finding conceptually refreshes cancer systems mechanism in which noncoding RNAs drive cancers, instead of proteins as conventionally thought[26–28]. This presents a novel basis for understanding the cancerous fundamental.

Pseudogenes were once thought as junk DNAs but recently they have been reported as regulators for cognate genes, in which they might regulate their corresponding protein-coding genes[8]. For example, pseudo-gene PTENP1 regulates PTEN in cancer. However, the number of known functional pseudogenes are very limited and the functions of these pseudo-genes have been thought as secondary. Here, we systematically revealed that the abundant pseudogenes were activated in cancer and these pseudogenes functionally worked as the most important cancer drivers instead of secondary regulators as thought. This was validated by clinical data in our paralleled study[16]. In contrast to the conventional validation via biochemistry in vitro in which results might not be applied to in vivo regulations, we systematically validated these noncoding RNA regulations by clinical data as in vivo evidences[16], which ensures our results more reliable than in vitro results.

Pseudogenes rarely target their cognate genes, but they mostly regulate their remote targets outside the chromosomes. Pseudogenes should execute their functions in a way similar to proteins as trans regulators and drivers. This further suggested that pseudogenes might act as flexible and energy-saving activators for various physiologic conditions. This opens the block around pseudo-genes to explore their functions in other physiologic conditions like stress stimulation.

Understanding the majority of noncoding RNAs working as cis-versus trans-regulators provides the first step to understand their functions and mechanisms, but it remains controversial due to lack of knowing the complete crosstalk involved in all noncoding

RNAs[8,9,11]. Here, we revealed that different types of noncoding RNAs have their own target-distance patterns varying with physiologic states, but universally, the majority of noncoding RNAs works as trans in normal, even antisense RNAs have only ~50% working in local (<1M). This parallels with the recent observation showing trans-regulation patterns in noncoding RNAs[29]. However, in cancer the majority of noncoding RNAs such as antisense RNAs and lincRNAs turns to target the local genes (<1Mb) as cis-regulators but not their cognates. Only a very limited number of noncoding RNAs target their cognates. Therefore, the hypothesized mechanism of noncoding RNAs executing their functions via bindings to complementary sequences of their cognates is misleading. In general, normal noncoding and coding genes primarily work as trans-regulators, but cancerous noncoding RNAs primarily serve as cis-regulators but not cognate-regulators.

Gene regulatory networks have been widely studied, but most of them have been derived from gene pair studies and condition-dependent experiments[8,11]. In addition, the current network inference approaches have suffered high noises and recently increasing noncoding RNA species have complicated the network inferences[8,9,11,13,14], resulting in seriously biased observations and leaving an actual blackbox of gene crosstalk. Here, we developed software to reveal the all endogenous crosstalk as systems networks hidden in massive data. Without any presumption, we generated the unbiased quantitative patterns from systems networks and revealed the systems mechanisms from the data patterns, which made our results reliable. To ensure our networks were robust, we only included interactions with high precision. High precision selections dramatically reduced the false positives and all interactions in our networks do not depend on any conditions. Obviously, some conventional interactions might not be found in our network due to they are conditional-dependent, not endogenous. Indeed we intentiontally missed numerous interactions that were conditionally dependent because including those condition-dependent interactions could dramatically introduce noise[14]. This practice to filter out noise to ensure reproducibility is also of first most concern in experimental biology, in which biologists normally conduct many experiments to prove true gene regulation. Here our computational algorithm has systematically revealed thousands of reliable regulations in two systems networks. These networks are invaluable and provide a novel foundation to advance our insights into cancer and human normal physiology.

## Declaration section

### Acknowledgments

Thank Dr Paul J Rider for reading and editing this manuscript.

### Availability of data and materials

All data resources and detailed network results were available on our project website (https://combai.org/network/)

### Authors’ Contributions

A.W. designed project, developed algorithm, coded software, and wrote the manuscript. R.H. was involved in project design and editing manuscript

